# Interleukins 4 and 13 Induce Exon Skipping of Mutant *Dystrophin* Pre-mRNA to Restore Dystrophin Production

**DOI:** 10.1101/207480

**Authors:** SiewHui Low, Chen-Ming Fan

## Abstract

Duchenne muscular dystrophy is (DMD) a lethal muscle degenerative disease caused by nonsense or out of frame deletion mutations in the *DMD* gene^1^, which encodes Dystrophin^2,3^. While multiple therapeutic strategies to ameliorate the disease symptoms are under development, there is currently no cure. Here we report an unexpected finding that intramuscular injections of the anti-inflammatory interleukin 4 or 13 (IL4/13) not only reduce inflammation but also restore Dystrophin protein production in the *mdx* mouse model^4^. IL4/13 restores Dystrophin production by inducing changes in the *Dmd* pre-mRNA splicing pattern that exclude the mutated exon and restore the reading frame. We further show that systemic delivery of IL4-Fc can restore Dystrophin in multiple muscle groups and increase muscle endurance and strength in *mdx* mice. Importantly, IL4/13 treatment of *mdx* myoblasts is sufficient to induce exon skipping and restore *Dmd* reading frame *in vitro*. Moreover, IL4-treated DMD patient myoblasts produce Dystrophin-positive myofibers after transplantation. In light of the established clinical safety of IL4 treatment^5,6^, we recommend IL4 as an agent of immediate consideration for treating Duchenne muscular dystrophy.

---

DMD is a devastating genetic disorder that starts with muscle weakness in early childhood and is ultimately lethal due to progressive muscle loss^1^. In skeletal muscles, the major *DMD* isoform consists of 79 exons and encodes a 427 kD Dystrophin protein^2,3^. Dystrophin is a structural protein that links the actin cytoskeleton to the sarcolemma and extracellular matrix to protect muscle against degeneration^7^. The *mdx* mouse carries a nonsense mutation in exon 23 (e23) of the orthologous *Dmd* gene and has been an invaluable a model for studying DMD^4^. This model also serves as a benchmark for evaluating pre-translational studies^8^. Currently, the most promising pre-translational approach is using the CRISPR technology to delete the mutated exon to restore reading frame^9,10^. However, CRISPR mediated therapies will likely require years of evaluation for safety due to potentially unexpected off target effects among highly heterogeneous patient genomes. Therefore, improving methods currently under development and finding new strategies remain of high priority^11^.

Because exons in the middle of the *DMD* locus are largely dispensable^12^, current methods are aimed to have internally truncated Dystrophin expressed via various means^13,14^. An estimated 80-90% of DMD patients can benefit from single or multiple exon skipping events to restore the reading frame and express internally truncated Dsytrophin^15^. Based on this, the use splice-site blocking anti-sense oligonucleotides to force exclusion of mutated exon to restore reading frame is in clinical trials^16^. Disease progression of DMD is also accompanied by chronic inflammation, and various anti-inflammatory treatments have been used to improve muscle pathology^17^. IL4 and its closely related IL13 are anti-inflammatory interleukins^18^ and both have been shown to enhance myoblast fusion *in vitro*^19,20^. IL4 also acts through fibro-adipogenic progenitors to promote muscle regeneration *in vivo*^21^. We therefore hypothesize that IL4 or IL13 treatment should have dual benefits of suppressing inflammation and enhancing muscle regeneration in the *mdx* mouse.

To examine the effect of IL4 and IL13 in the *mdx* mouse, we injected each into the tibialis anterior (TA) muscles of the right leg, and PBS into the left TA muscles of the same animals as controls (Fig. 1a). Because *mdx* mice develop pathology at ~3 weeks after birth^4^, we chose early intervention at week 2 after birth (2 μg of IL4 or IL13 per kg body weight (2 μg/kg) per TA muscle). Consistent with their anti-inflammatory role, IL4-and IL13-treated muscles displayed fewer interstitial cells and infiltrating F4/80-positive (F4/80^+^) macrophages, compared to control muscles (Fig. 1b-d). Unexpectedly, when we examined Dystrophin expression using the MANDRA1 antibody against its C-terminus (a.a. 3667-3671, encoded in exon 77)^ref.22^, we found that IL4- and IL13-treated *mdx* TA muscles had significantly more MANDRA1^+^ myofibers (>20%) than control *mdx* muscles (0.6%; Fig. 1e,f). Western blot analyses confirmed increased levels of Dystrophin in IL4- and IL13-treated *mdx* muscle lysates (Fig. 1g,h). Using Evans Blue Dye (EBD) to assess membrane permeability, we determined that IL4- and IL13-treated samples had reduced EBD^+^ muscle fibers, reflecting an improved myofiber integrity (Fig. 1i,j). Pharmacokinetic assessment of IL4 and IL13 (Supplementary Fig. S1a,b) revealed the strongest effects (based on MANDRA1^+^ myofibers) of both ILs was at 2 μg/kg in the 2- and 3-week age groups, while a decreased effect at 5 μg/kg was noted. Treatment proved less effective as mice matured (weeks 4-10). Thus early intervention by IL4 and IL13 can reduce inflammatory infiltration and restore Dystrophin in *mdx* TA muscles.

**Fig. 1.**
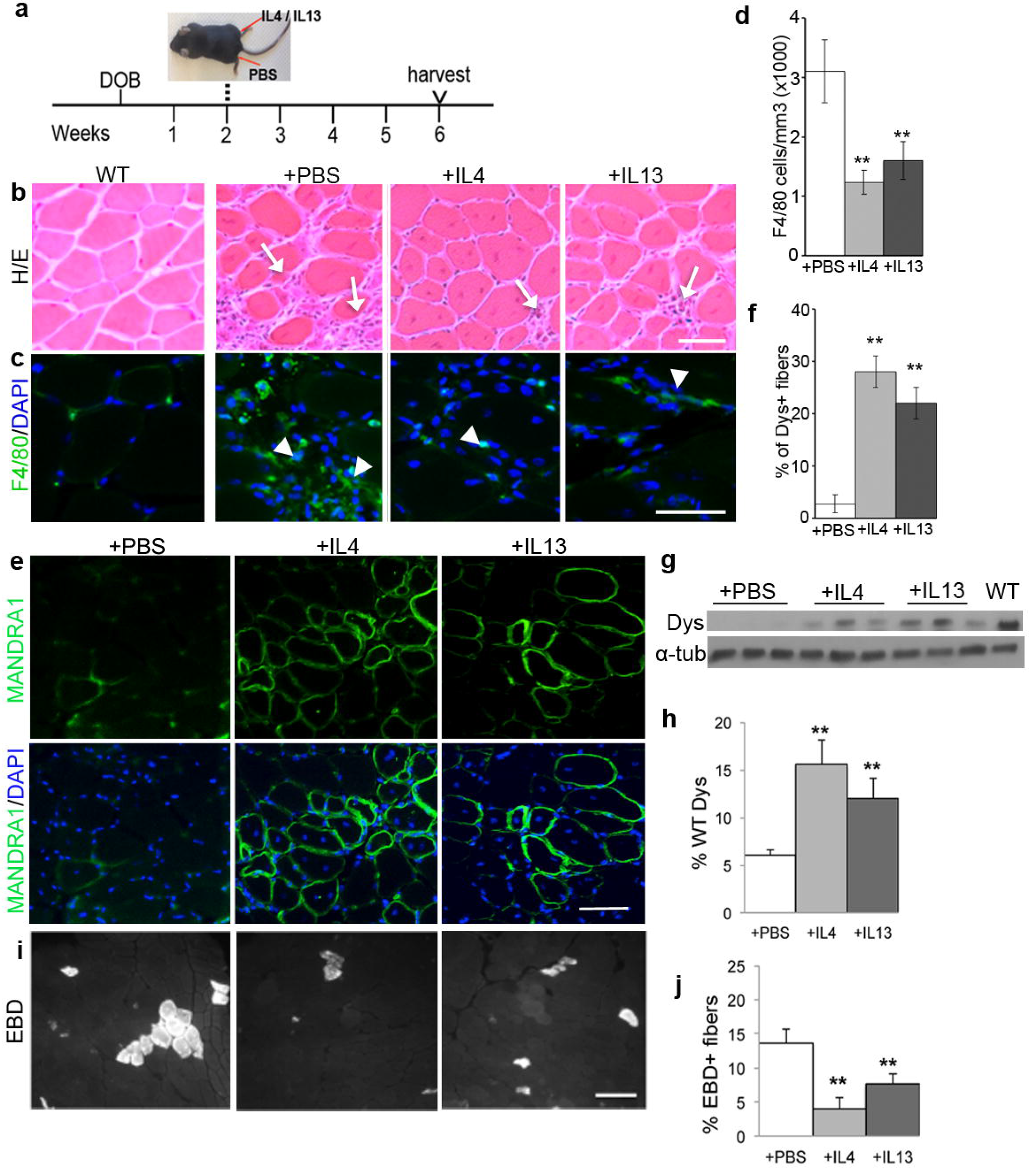
IL4 and IL13 improve *mdx* pathology and restore Dystrophin expression. **(a)**Diagram of the assay scheme: TA muscles of 2-week old *mdx* mice were injected with IL4 or IL13 (right TA) and PBS (left TA) and harvested 4 weeks later; DOB, day of birth. **(b)** Hematoxylin and eosin (H/E) stained sections of wild type (WT), and *mdx* TA muscles treated with PBS (+PBS), IL4 (+IL4), and IL13 (+IL13); arrows, fibrotic areas. **(c)** Immunostaining by F4/80 (green) for infiltrating macrophages (arrowheads); nuclei stained with DAPI (blue). **(d)** Quantification of data in **(c)**. **(e)** Immunostaining by MANDRA1 for Dystrophin (green) of PBS-, IL4-, and IL13-treated *mdx* TA muscles (top); nuclei stained with DAPI (blue) and merged images at the bottom. **(f)** Percentage of Dys^+^ (MANDRA1^+^) fibers from data in **(e)**. **(g)** Western blot for Dystrophin (Dys) of PBS-, IL4-and IL13-treated *mdx* muscles; wild type (WT), for comparison; α-tubulin (α-tub), loading control. **(h)** Quantification of data in **(g)**, normalized to α-tub. **(i)** EBD staining in PBS-, IL4- and IL13-treated *mdx* TA muscles. **(j)** Percentage of EBD_+_ fibers in **(i)**. Bar graphs are shown as mean ± SEM (n = 3 for each group); ** p ≤ 0.001. Scale bars = 50 μm.

Using RT-PCR, we found multiple spliced forms that exclude the mutated exon and predict contiguous reading frame in IL4/13-treated *mdx* muscles (Fig. 2a), indicating that skipping of the mutated exon 23 underlies Dystrophin restoration. Given that the *DMD* transcript has a half-life of ~16 h^23^ and that exon-skipped *Dmd* transcripts are found weeks after IL4/13 injection, these exon-skipping events appear persistent. Multiple splicing variants predict multiple Dystrophin isoforms. To examine this, we used *Dmd* exon-specific antibodies MANDYS18 (to an epitope encoded in e26) and MANEX4850A (to an epitope encoded in e48-50)^ref.22^ together with MANDRA1. Double staining of MANDYS18 and MANDRA1 showed mosaic distribution of these epitopes (Fig. 2b-d), reflecting that induced Dystrophin^+^ fibers express different variants. Similar results were found for MANEX4850A and MANDRA1 double staining (Supplementary Fig. S2). Accompanying Dystrophin production, Dystrobrevin and Syntrophin were also restored at the sarcolemma of treated *mdx* muscles (Fig. 2e-g), suggesting formation of functional complex. Thus, IL4 and IL13 induce exon skipping to restore Dystrophin and its associated proteins to the *mdx* sarcolemma.

**Fig. 2.**
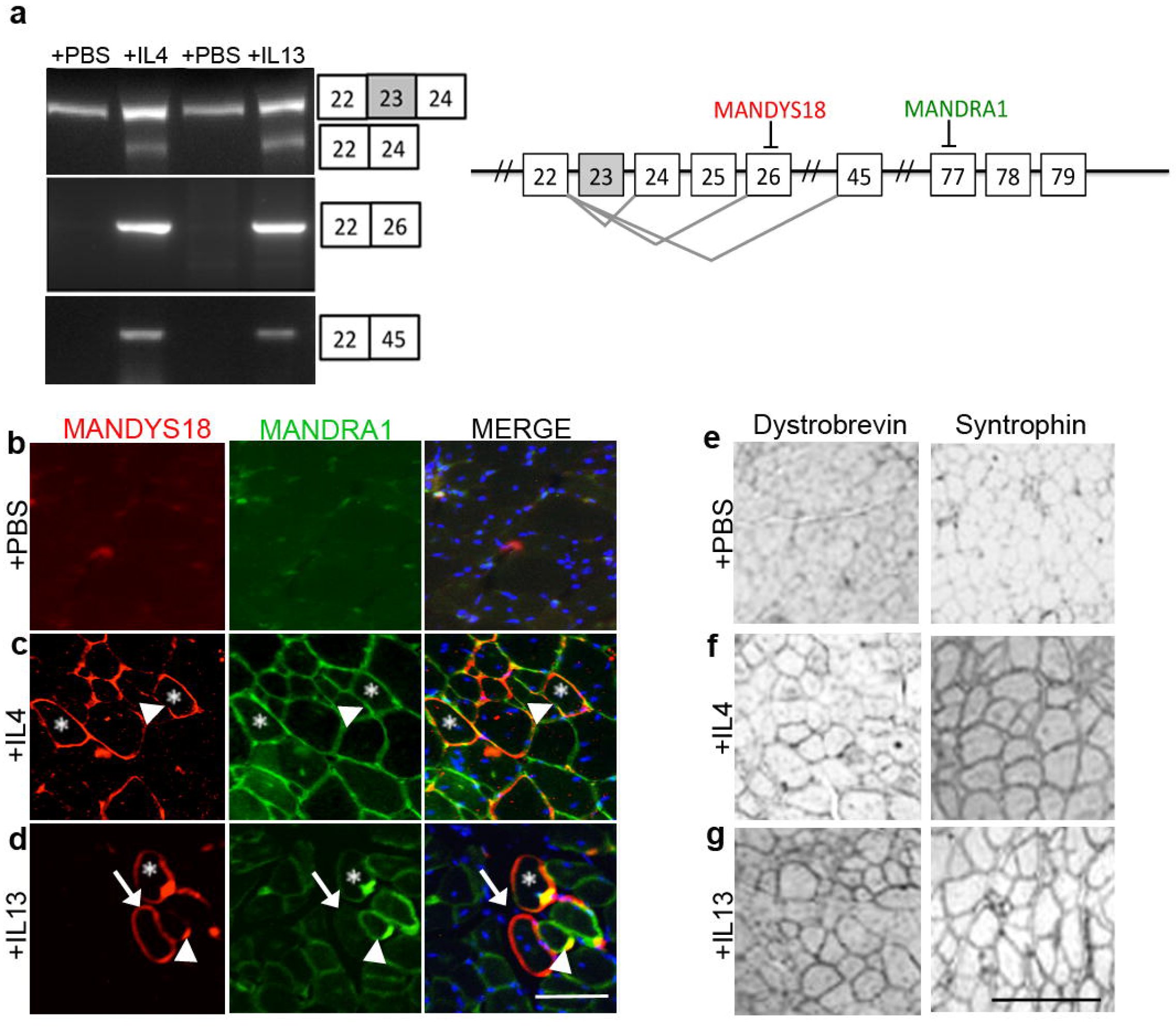
IL4 and IL13 induce multiple in-frame exon-skipping events and restore Dystrophin and its associated proteins. **(a)** RT-PCR detected e22-24, e22-26, and e22-45 joined transcripts induced by IL4 and IL13, relative to PBS-treated TA muscles; nonskipped e22-23-24, input control. Splicing patterns are summarized at the right, with the locations of epitopes recognized by MANDYS18 and MANDRA1. **(b-d)** Examples of double-immunostaining by MANDYS18 (red) and MANDRA1 (green) of **(b)** PBS-, **(c)**IL4-, and **(d)** IL13-treated *mdx* muscles; nuclei stained with DAPI (blue); merged images to the right. Asterisks indicate MANDRA1^+^MANDYS18^+^ myofibers; arrowheads, MANDYS18^−^MANDRA1^+^ myofibers; arrows, MANDYS18^+^MANDRA1^−^ myofibers. Note the uneven distribution of epitopes within a myofiber. Scale bar = 50 μm. **(e)** Immunohistochemistry of PBS, IL4-, and IL13-treated *mdx* TA muscles with antibodies specific for Dystrobrevin and Syntrophin. Scale bar = 200 μm.

Transcriptomic analyses revealed 200 up-regulated and 254 down-regulated genes common in IL4- and IL13-treated *mdx* muscles, compared to PBS-treated *mdx* muscles (Supplementary Fig. S3). Forty-eight of these genes showed a reversed trend of expression from that in DMD patients and animal models^24–28^, supporting an improvement. As expected from known actions of IL4 and IL13, a larger fraction of skeletal muscle tissue/fiber developmental genes shows upregulation, while an increased fraction of inflammatory genes is repressed.

Systemic delivery of IL2-Fc was recently reported to ameliorate histopathology of *mdx* muscles^29^. We were inspired to test whether systemic IL4-Fc could restore Dystrophin in different muscle groups. IL4-Fc and IL2-Fc (at 10 μg/kg bodyweight) were injected peritoneally into 2 weeks old *mdx* mice for comparison. TA, diaphragm and gastrocnemius muscles were analyzed 4 weeks later. IL4-Fc treated *mdx* mice contained significantly more MANDRA1^+^ myofibers in all 3 muscle groups, compared to PBS- and IL2-Fc-treated groups (Fig. 3a-d). Western blot analyses confirmed increased Dystrophin levels (Fig. 3e-h) and EBD staining confirmed increased membrane integrity (Supplementary Fig. S4) in these muscle groups. To determine whether muscle strength and resistance were improved, we assessed forelimb weightlifting capability (Fig. 3i) and used the Kondziela’s inverted test for muscle resistance and strength (Fig. 3j)^30,31^. As expected, IL4-Fc treated group performed better on both tests than the PBS-treated group.

**Fig. 3.**
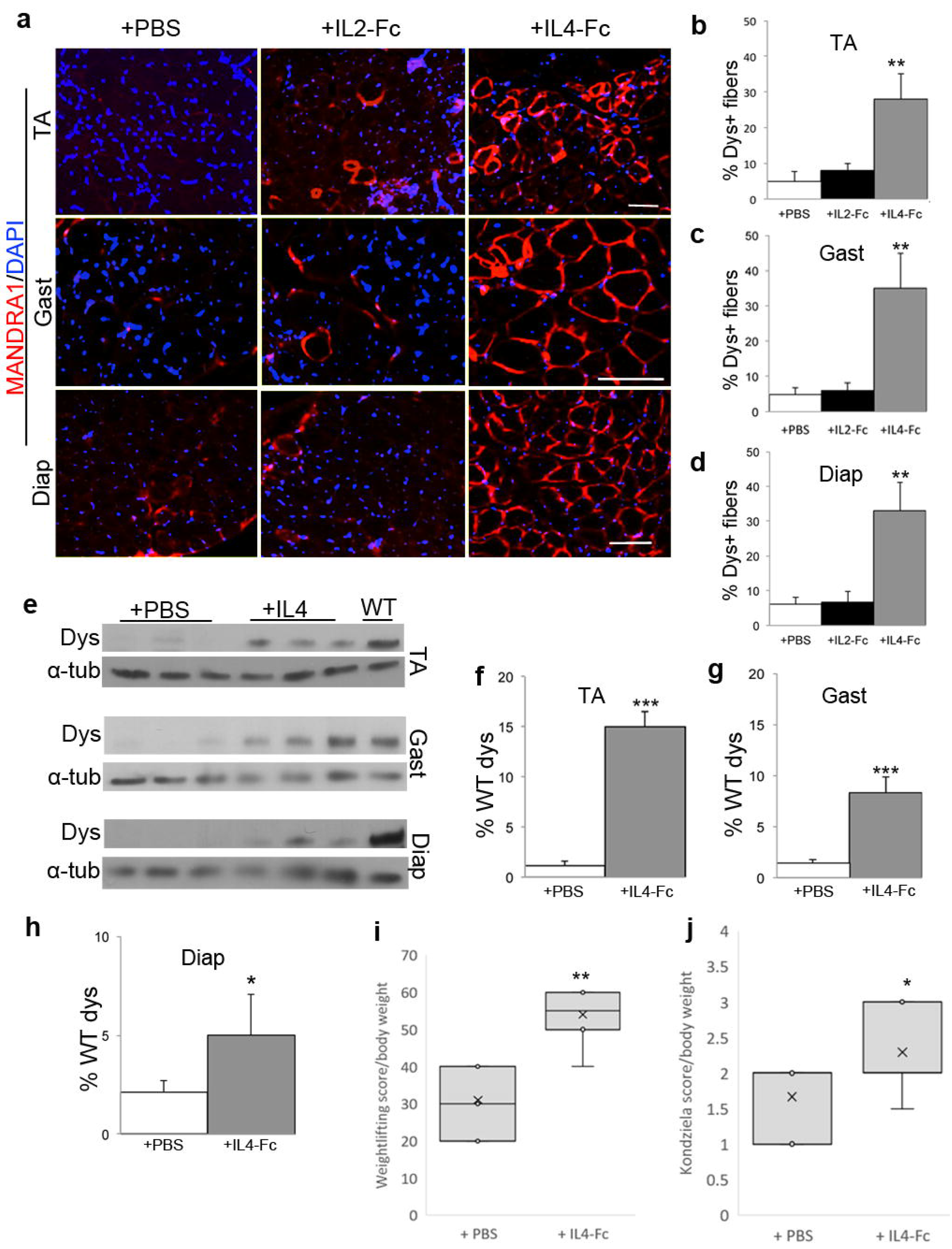
Intraperitoneal injection of IL4-Fc restores Dystrophin and improves muscle strength. 2-week old *mdx* mice were injected with PBS, IL2-Fc or IL4-Fc intraperitoneally, and various muscle groups were harvested 4 weeks later. **(a)** Immunostaining by MANDRA1 (red) of TA, Gastrocnemius (Gast) and Diaphragm (Diap) muscles from *mdx* mice injected with PBS, IL2-Fc, and IL4-Fc; nuclei stained with DAPI (blue). **(b-d)** Percentage of Dys^+^ fibers in a section from **(b)** TA, **(c)** Gast, and **(d)** Diap, from data in **(a)**. **(e)** Western blots for Dys in TA, Gast, and Diap muscles from PBS-and IL4-Fc-treated *mdx* mice; α-tubulin (α-tub), loading control. Three biological replicates of each group are shown, and the WT sample, for comparison. **(f-h)** Quantification of Dys levels from **(f)** TA, **(g)** Gast, and **(h)** Diap muscles, normalized to α-tub and presented as percentages of the WT level. **(i)** Weightlifting test of forelimbs muscle strength and **(j)** Kondziela’s inverted test for muscle strength and resistance using all four limbs, for PBS-and IL4-Fc-treated *mdx* mice. The scores were normalized by the body weight. Bar graphs are shown as mean ± SEM (n =3 for each group); ** p ≤ 0.001

We next examined whether IL4 and IL13 acted directly on *mdx* myoblasts to induce *Dmd* exon skipping. Using a modified myoblast recruitment protocol^19^, we isolated myoblasts from ~2 weeks old *mdx* mice and treated them with IL4 or IL13 for 48 h in growth media. These cells were then exposed to differentiation media for 72 h and assayed for Dystrophin expression (Supplementary Fig. S5a). Treated *mdx* myoblasts gave rise to ~50% MANDRA1^+^ cells, while control *mdx* cells showed ~5-10% spontaneously reverted cells (Fig. 4a,b). After normalization to Myogenin (MyoG)^+^ nuclear number (Fig. 4c), the MANDRA1-inducing effect of IL4/13 remained. This result was unlikely a consequence of increased *mdx* myoblast proliferation as IL4/IL13 displayed only a modest mitogenic activity (Supplementary Fig. S5b). Furthermore, e22-24 joined *Dmd* transcripts were detected by RT-PCR in IL4/IL13-treated myoblasts (Fig. 4d). Titration data indicated that Dystrophin restoration activity of IL4 and IL13 plateaued at 5 and 10 ng/ml, respectively, without a negative impact up to 100 ng/ml (Supplementary Fig. S5c), contrasting to their narrow effective dose *in vivo*. We also confirmed in our culture the absence of the PDGFRα fibro-adipogenic progenitor (Supplementary Fig. S5d), an IL4 target cell type that indirectly enhances muscle regeneration^21^.

**Fig. 4.**
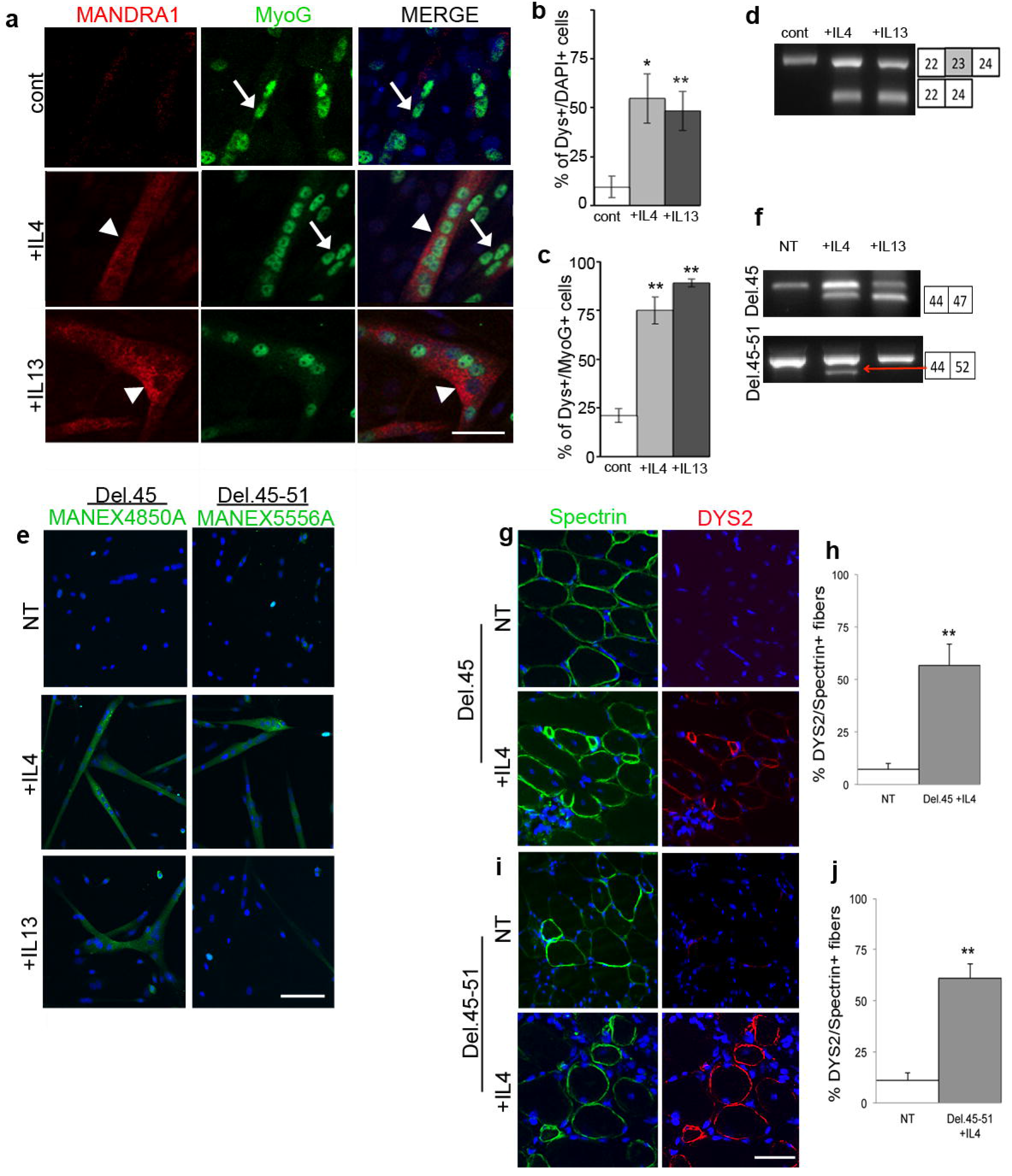
IL4 restores Dystrophin expression in *mdx* and DMD myoblasts in cultures and after transplantation. **(a)** Primary *mdx* myoblasts were mock-treated (cont) or treated with IL4 or IL13 (See Fig. S5a for assay scheme) and immunostained for MANDRA1 (red) and MyoG (green) of PBS-, IL4- and IL13-treated *mdx* cultures; nuclei stained with DAPI (blue); merged images to the right; arrowheads, MANDRA1^+^MyoG^+^ cells. Scale bar = 30 μm. **(b)** Percentages of MANDRA1^+^ (Dys^+^) overlapping signals with DAPI^+^ nuclei from data in **(a)**. **(c)** Percentages of MANDRA1^+^ (Dys^+^) signals overlapping with MyoG^+^ nuclei from data in **(a)**. For **(b, c)**, bar graphs are mean ± SEM (n =3 for each); * *p* ≤ 0.01, \*\**p* ≤ 0.001. **(d)** RT–PCR detected e22-24 joined transcripts induced by IL4 and IL13; un-skipped e22-23-24, input control. **(e)** Two human DMD myoblast lines with different exon deletions (Del.45, deletion of exon 45; Del.45-51, deletion of exons 45-51) were cultured as mentioned in **(a)** and immunostaining by MANEX4850A (green) for the Del.45 line, and MANEX5556A for the Del.45-51 line from non-treated (NT), IL4- and IL13-treated samples; nuclei stained with DAPI (blue). Scale bar = 50 p,m. **(f)** RT-PCR detected e44-46 and e44-52 joined transcripts induced by IL4 or IL13 in the two DMD myoblast lines. **(g-j)** IL4-treated **(g)** Del.45 and **(i)** Del.45-51 myoblasts transplanted into the TA muscle of NRG mice were harvested 4 weeks later and immunostained for human-specific Spectrin (green) and DYS2 (red); nuclei stained with DAPI (blue); merged images to the right. Percentages of DYS2 in Spectrin+ cells from data in **(g)** and **(i)** are in **(h)** and **(j)**, respectively; bar graphs are shown as mean ± SEM (n = 3 for each group); ** p ≤ 0.001. Scale bars = 50 μm.

qRT-PCR profiling of key signaling components *of* IL4 and IL13 revealed that (except for *Tyk2*) *IL4Rα*, *Jak1*, *Jak2*, *Jak3*, and *Stat6* are expressed at higher levels in 2-week than in 3-mon old *mdx* myoblasts (Supplementary Fig. S6a), providing a possible explanation for the age-dependent response *in vivo*. Importantly, Dystrophin restoration in *mdx* myoblasts induced by IL4 is dependent on IL4Ra, as this effect was abolished when we performed siRNA knockdown of *IL4Ra*, compared to a control scrambled siRNA (*siScr*) (Supplementary Fig. S6b-d). We also found that inhibition of JAK1 and JAK2 by pharmacological inhibitors negates IL4-induced Dystrophin reversion (Supplementary Fig. S6e,f), providing additional support for a direct effect of IL4 signaling in myoblast. We further conducted RNA-seq of *mdx* and IL4-treated *mdx* samples and analyzed their splicing patterns. IL4 did not cause a significant change in the overall mRNA processing pattern (Supplementary Fig. S7), suggesting that exon-skipping of the mutant *Dmd* locus is not the consequence of a global splicing change.

As we applied IL4 to *mdx* cells at the myoblast growth stage and observed Dystrophin reversion in differentiated myotubes, we asked whether IL4 treatment could have a long-term effect. For this, we obtained reserve cells^32^, similar to quiescent muscle stem cells, from differentiated *mdx* cultures that were initially un-treated or treated with IL4 (Supplementary Fig. S8a,b). We then re-activated them to growing myoblasts and differentiated them again, all the while without adding IL4. Intriguingly, IL4-treated *mdx* myoblasts retained Dystrophin reversion phenotype in the second round of differentiation (Supplementary Fig. S8c,d). This predicts that IL4/13 treatment *in vivo* may also maintain myofibers with restored Dystrophin over a long period. Indeed, 30 weeks after a single intramuscular injection of IL4 or IL13, considerable MANDRA1+ myofibers were found (Supplementary Fig. S8e-g). Altogether, we conclude that IL4, and likely IL13, acts directly on *mdx* myoblasts to induce a long-term Dystrophin reversion phenotype.

To begin evaluating the viability of IL4/13 as a treatment for DMD, we explored whether the exon skipping/Dystrophin restoration activity of IL4 and IL13 was effective on DMD patient-derived immortalized cell lines^33^. We chose two lines that were derived from juveniles and represented frequently deleted genomic region of *DMD*, Del.45 and Del.45-51. They were treated with IL4 or IL13, induced to differentiated, and assayed for Dystrophin. For the Del.45 line, MANEX4850A^22^ were used for assessment (Fig. 4e), and for the Del.45-51 line, MANEX5556A^22^ (to epitopes encoded in exons 55-56; Fig. 4e). These epitopes were not detected in non-treated (NT) cell lines, but present after treatment of IL4. Curiously, IL13 induced the MANEX4850A epitope in the Del.e45 line, but not the MANEX5556A epitope in the Del.45-51 line. RT-PCR data confirmed e44-46 and e44-52 joined transcripts in IL4-treated Del.45 and Del.45-51 lines, respectively, and e44-46 joined transcripts in IL13-treated Del.45 line (Fig. 4f).

Finally, we tested whether xenotransplantation of IL4-treated Del.45 and Del.45-51 cells into cryodamaged TA muscles of NRG mouse would yield Dystrophin^+^ myofibers. One month after transplantation, we found human-specific Spectrin^+^ myofibers derived from transplanted Del.45 (Fig. 4g) and Del.45-51 cells (Fig. 4i) regardless of treatments. Within the Spectrin^+^ area, significantly more DYS2^+^ myofibers were found in IL4-treated, compared to non-treated Del.45 (Fig.4g, h) and Del.45-51 (Fig.4i, j) cells. Thus, IL4-treated DMD cell lines show restored Dystrophin expression *in vivo*.

Our study defines a new role for IL4 and IL13 and expands their functional repertories beyond immune responses^18^, myoblast recruitment/muscle regeneration^18–20^, and brown adipogenesis^34^. Interestingly, we found IL4/13 treatment most effective on juvenile animals and myoblasts. We propose IL4/13 utilizes a developmental window to affect long-term changes in pre-mRNA splicing pattern. As IL4 induces exon-skipping events around the middle exons of the *Dmd/DMD* loci, it may have a wide-range of application to a larger cohort of patients than the specific designs needed for anti-sense oligos or CRISPR to each patient. In contrast to current strategies employing transgenes, stem cells, oligonucleotides, and CRISPR, IL4 and IL13 are natural biomolecules and the safety of IL4 has been established in clinical trials^5,6^. How IL4/IL13 induces exon skipping of the mutated *Dmd* locus in mouse and human patient myoblasts remains an intriguing open question to be answered. We therefore urge clinical teams specialized in DMD to consider fast track the process and begin trials of applying our approach to patients.

## Methods

(Additional protocols and reagents are detailed in supplementary information).

### Animals and Treatments

C57BL/10, *mdx* and NRG mice (Jackson Laboratory) were housed in a pathogen-free facility. All animal experiments were approved by IACUC. PBS, IL4 and IL13 (R&D Systems) were delivered intramuscularly. IL2-Fc, IL4-Fc (AdipoGen Life Sciences), and Evans Blue dye (EBD, Sigma) were delivered intraperitoneally. NRG mice were used as transplantation recipients for human DMD myoblasts. Age of animals and muscle sample harvest time line are specified in the text and legends.

### Processing of muscle samples

Muscle samples were cryosectioned at 8 μm. For histology or immunofluorescence, sections were fixed by 4% paraformaldehyde (in PBS), and subjected to hematoxilin & eosin (Surgipath) staining or primary and secondary antibody incubation, respectively, as described^35^. For visualizing EBD staining, sections were fixed in cold acetone (−20°C). Antibody sources and dilutions are detailed in supplementary information.

### Western blotting

Muscle protein extracts were prepared as described^35^ and resolved by SDS-PAGE (4-20% gradient gel, Bio-Rad), followed by Western blotting using specified antibodies and ECL detection (GE Healthcare).

### Primary myoblast culture

Primary myoblasts were isolated as described^18^ and cultured on Matrigel (BD Biosciences) coated dishes or chamber slides. PBS, IL4 or IL13 was added directly to growth media. Cultures were then washed and incubated for 72 h in differentiation medium, and used for immunostaining, protein extracts, RT-PCR or RNA-seq. Reserve cells were isolated by partial trypsinisation^32^ of 4-day differentiated cultures. For siRNA (Life Technologies) transfection, Lipofectamine 3000 (Invitrogen) were used. JAK1-3 inhibitors (Calbiochem) were applied 6 h prior to IL4 addition and throughout culture period. For myoblast proliferation, Cell Titer 96 Aqueous One Solution Cell Proliferation Assay kit (Promega) was used.

### RNA extraction, RT-PCR, nested RT-PCR and quantitative PCR (qPCR)

Total RNAs from TA samples and cultured cells were extracted using TRIzol reagent (Invitrogen) and used for RT-PCR, qRT-PCR, or RNA-seq. Exon skipped *Dmd* transcripts were detected using nested primers to detect in-frame splicing events. PCR products were resolved by electrophoresis on a 2% agarose gel. qPCR was performed using the BioRad CFX96 Real-Time System with CFX manager.

### Muscle strength

Kondziela’s test and a weightlifting test were performed to evaluate strength, resistance, and exercise abilities^30, 31^. The scores were calculated and normalized by the body weight.

### Human DMD immortalized cell culture and transplantation

Human DMD cell lines were maintained in Skeletal Muscle Cell Growth Medium (Promocell). IL4 and IL13 (R&D Systems) treatment was the same as for *mdx* myoblasts. For transplantation, 5×10^5^ cells were used for injection into TA muscles of NRG immunodeficient mice^33^.

### RNA sequencing and bioinformatics

For each RNA-seq data set, 3 independent samples were used. Purity and integrity of RNA were assessed by nanodrop ND-100 (Nanodrop) and Bioanalyser 2100 (Agilent). cDNA libraries were constructed using Illumina TruSeq RNA Library Prep Kit v2, and sequenced by HiSeq2000; 100-nucleotide single read and ~40 million reads per sample. Normalized read counts (in FPKM) were calculated using Bowtie2 v 2.0.6, TopHat v 2.0.7, and Cufflinks version 2.02. Functional analysis was done by DAVID functional annotation clustering and for up-regulated and down-regulated genes with a fold change of ≥ 2 and enrichment score of ≥ 1. Quantas pipeline was used for analysis of alternative splicing information from RNA-Seq data.

### Image Acquisition and Processing

All images were taken by AxioCam mounted on an Axioscope. Signal intensities were adjusted only when necessary for presentation according to the ORI data processing guidelines. No quantification statements were made based on difference in fluorescence intensity.

### Quantification and statistical Analyses

Results are expressed as means ± SEMs, unless otherwise specified. Statistical analyses were performed by Excel using *t*-test. A *p*-value less than 0.05 is considered significant.

## Accession codes

All sequencing data have been deposited at NCBI Sequence Read Archive (SRA), with assigned project number PRJNA414242 (to be released).

## Acknowledgements

We thank E. Dikovskaia and S. Satchell for technical assistance, A. Pinder and F. Tan for RNA-seq, and M. Sieber, S. Southard, and Fan lab members for comments. We especially thank Dr. V. Mouly for DMD immortal cell lines and Dr. Glenn Morris for sharing the antibodies. The Carnegie Institution for Science and National Institute of Arthritis and Muscular and Skin diseases of the National Institutes of Health (NIH) (AR060042 to C.-M.F.) provided funding for this work. S.L. is supported by the same NIH grant.

## Author Contributions

S.L. and C.-M.F. conceptualized the study. S.L. performed experimental analysis. S.L. and C.-M. Fan planned, wrote, discussed and edited the manuscript.

